# Directed evolution of bacteriophages: impacts of prolific prophage

**DOI:** 10.1101/2024.06.28.601269

**Authors:** Tracey Lee Peters, Jacob Schow, Emma Spencer, JT Van Leuven, Holly Wichman, Craig Miller

## Abstract

Various directed evolution methods exist that seek to procure bacteriophages with expanded host ranges, typically targeting phage-resistant or non-permissive bacterial hosts. The general premise of these methods is to propagate phage on multiple bacterial hosts, pool the lysate, and repeat the propagation process until phage(s) can form plaques on the target host(s). In theory, this propagation process produces a phage lysate that contains input phages and their evolved phage progeny. However, in practice, this phage lysate can also include prophages originating from bacterial hosts. Here we describe our experience implementing one directed evolution method, the Appelmans protocol, to study phage evolution in the *Pseudomonas aeruginosa* phage-host system, in which we observed rapid host-range expansion of the phage cocktail. Further experimentation and sequencing analysis revealed that this observed host-range expansion was due to a *Casadabanvirus* prophage that originated from one of the Appelmans hosts. Host-range analysis of the prophage showed that it could infect five of eight bacterial hosts initially used, allowing it to proliferate and persist through the end of the experiment. This prophage was represented in half of the sequenced phage samples isolated from the Appelmans experiment. This work highlights the impact of prophages in directed evolution experiments and the importance of incorporating sequencing data in analyses to verify output phages, particularly for those attempting to procure phages intended for phage therapy applications. This study also notes the usefulness of intraspecies antagonism assays between bacterial host strains to establish a baseline for inhibitory activity and determine presence of prophage.

**IMPORTANCE:** Directed evolution is a common strategy for evolving phages to expand host range, often targeting pathogenic strains of bacteria. In this study we investigated phage host-range expansion using directed evolution in the *Pseudomonas aeruginosa* system. We show that prophage are active players in directed evolution and can contribute to observation of host-range expansion. Since prophage are prevalent in bacterial hosts, particularly pathogenic strains of bacteria, and all directed evolution approaches involve iteratively propagating phage on one or more bacterial hosts, the presence of prophage in phage preparations is a factor that needs to be considered in experimental design and interpretation of results. These results highlight the importance of screening for prophages either genetically or through intraspecies antagonism assays during selection of bacterial strains and will contribute to improving experimental design of future directed evolution studies.

## INTRODUCTION

Directed evolution of phages is a term used to describe the experimental methods that promote genetic and phenotypic evolution of phages towards some desired trait. Directed evolution methods vary by host presentation strategy and typically fall into three categories: sequential, parallel or mixed(1). In all cases, the general strategy of directed evolution methods includes iteratively propagating phages on one or more hosts, then harvesting the output phage lysate after growth, and using it as the input phage(s) for the following round. Also referred to as phage training, one goal of directed evolution is to procure phages with altered or expanded host range targeting phage-resistant or otherwise non-permissive hosts(2).

The Appelmans protocol is one directed evolution method that is used to expand the host range of bacteriophages for phage therapy applications(2–4). This method employs a parallel presentation of bacterial hosts to a phage cocktail of two or more phages, where the phage mixture is serially diluted and allowed to propagate on individual hosts, followed by harvesting pooled lysate. The pooled phage lysate is then again diluted and allowed to propagate on individual hosts for a variable number of subsequent iterations(2, 3). The end goal is to obtain a phage lysate or an individual phage isolate that has evolved to infect a desired number of target bacterial hosts. Subsequent infections increase the likelihood of coinfection and recombination, thus increasing the potential for diversity in the resulting pooled phage lysate, without the addition of new genetic information(4).

In theory, as the input phage cocktail is propagated on the hosts, pooled lysate includes both input and evolved phage progeny. In practice, basic methodology for harvesting the pooled phage lysate in subsequent rounds results in the inclusion of input phages, their potentially evolved progeny, and inevitably, various cellular components and bacterial by-products, such as resident prophage induced from the bacterial host(s).

Many pathogenic bacterial strains of clinical importance carry prophage gene clusters within their genomes(5–7). Prophage gene clusters may represent cryptic prophages such as tailocins, filamentous phages, and active inducible prophages (8–11). Exposure of a prophage-carrying host (otherwise known as a lysogen) to mitomycin C or ultra-violet light can often lead to prophage induction, although some prophages may be produced at low levels (previous studies reported induction rates of 0.09% to 3.1% of the bacterial population) due to spontaneous DNA damage and heterogenous expression of genes involved in the SOS response(12, 13). Prophage may be maintained by the host because they can contribute some survival and fitness advantage such as promoting biofilm formation, bacterial virulence, immunity to subsequent phage infections, and horizontal gene transfer(14–16). Thus, the presence of prophage in bacterial hosts can be considered a source of additional “surprise” genetic input in directed evolution conditions.

Although the prevalence of prophages is well established and directed evolution methods are common practice, the phenomenon of prophage induction during experimental evolution of phages and the potential influence on experimental outcomes or data interpretation has been underreported until recently. For example, Vu et al. found that prophage induction and evolution occurred during the Appelmans protocol in the *Acinetobacter baumanni* phage-host system and contributed to expanded host range of the phage cocktail(17). Reporting and acknowledging the presence and influence of prophages in directed evolution experiments will contribute to improved experimental design for future studies.

Here we describe our experience employing the Appelmans protocol to study phage evolution on *Pseudomonas aeruginosa*, in which prophage induction early in the experiment was responsible for the observed rapid host-range expansion. The wildtype prophage had the ability to infect five target bacterial hosts, allowing it to proliferate and persist through the end of the protocol. This prophage remained at high abundance, and was represented in half of the phage samples isolated from the experiment, as confirmed by whole genome sequencing. This phage was found to be closely related to a known lysogenic, generalized transducing phage, JBD24 of the genus *Casadabanvirus*(18). This work highlights the importance of genetically verifying input and output phages in directed evolution studies and serves as a cautionary tale for future design of directed evolution experiments, particularly for those attempting to procure phages for phage therapy applications(19). We also outline the utility of intraspecies antagonism assays for identifying underlying cross-strain activity due to tailocins or resident prophages.

## RESULTS

### Using directed evolution to expand phage host range

The purpose of this work was to study genetic determinants of phage evolution in context of phage host-range expansion. The goal was to expand the host range of three input phages onto non-permissive hosts, then identify genetic mutations that may have contributed to the phenotypic change. Thus, we endeavored to implement the Appelmans protocol in *Pseudomonas aeruginosa,* a human pathogen of high research priority (see **Figure 1** for experimental design)(2). Three phages were used as the input phage cocktail: D3, M6, and JM2(20, 21). Five laboratory strains (three permissive, two non-permissive) and three clinical isolates (non-permissive) were originally used for the first three rounds of the protocol. Preliminary host-range analysis of the input phages established that they could each infect two hosts, PAO1 and either PA103 or PDO300, collectively infecting three of eight host strains initially selected. Phages were 10-fold serially diluted and added to the respective wells for each bacterial host. After overnight incubation, all cleared wells and the least dilute uncleared well for each host were pooled, and lysate prepared as input phage for the subsequent round.

**Figure 1.**
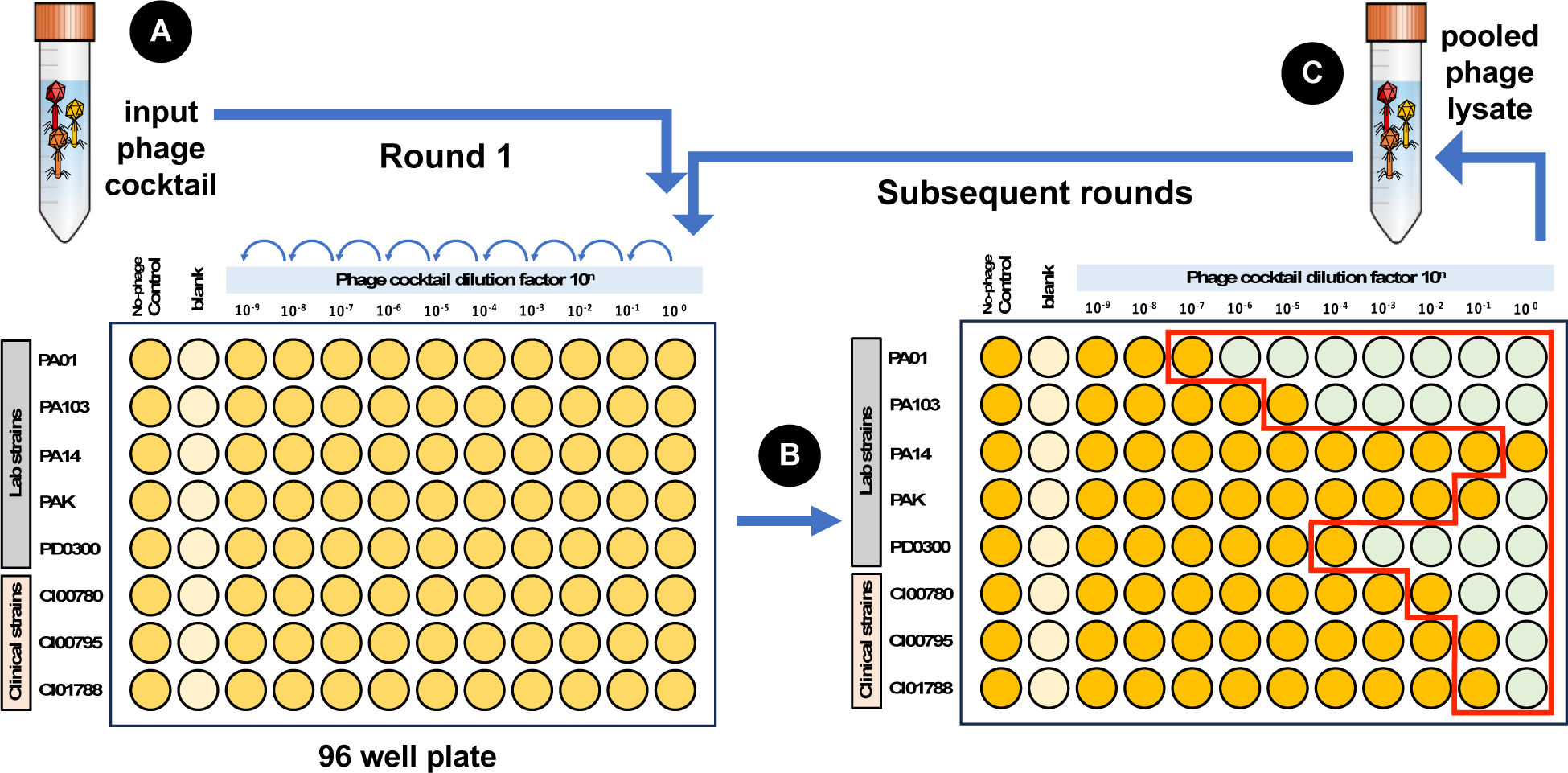
Experimental layout of Appelmans protocol for 96-well plate. (**A**) Input phage cocktail is diluted and applied to each bacterial host; (**B**) Overnight incubation results in cleared wells (green) and turbid wells (yellow); (**C**) Wells are pooled (red outline) and used as the input phage cocktail for subsequent rounds.

### Rapid host-range expansion and lysis of non-permissive hosts

The Appelmans protocol was implemented for a total of nine rounds. Surprisingly, host-range expansion of the phage cocktail was observed within the first four rounds of passaging, with clearing of non-permissive hosts PAK (in the second round) and PA14 (in the fourth round)(**Figure 2**). The three clinical isolates were removed from the experiment after the third round due to slow growth rates compared to the laboratory strains. Pooled phage lysate was plated onto lawns of target strains and phages were isolated by plaque purification on individual hosts. Host-range analysis of output phages showed that five phage isolates with expanded host range could form plaques on all five target hosts, PAO1, PA103, PDO300, PAK, and PA14 (**Figure 3**). Three of these phages were isolated after round three and two were isolated after round nine on either strain PAK or PA14. No plaques were observed on the clinical isolate strains. Ten output phages of interest were prepared for DNA extraction and submitted for sequencing. compared to the three input phage genomes, D3, M6 and JM2. Phages D3, M6 and JM2 have genomes of approximately 56kbp, 61kbp, and 60kbp respectively (**Supplemental Figure 2**). Surprisingly, assemblies for the five output phages with expanded host range (**Figure 3**: PAK_R3_01, PAK_R9_07, PA14_R3_02,

**Figure 2.**
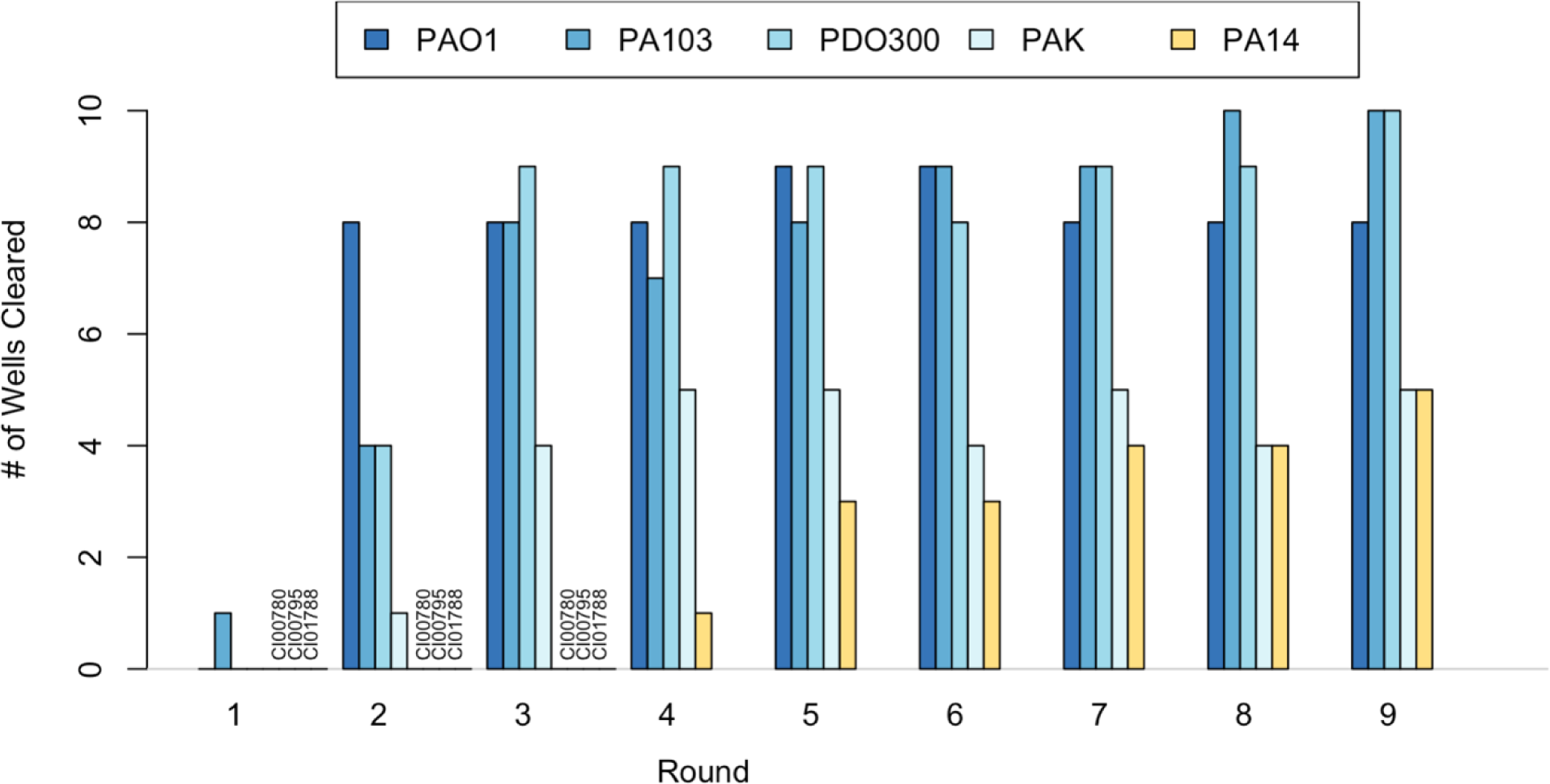
Directed evolution of phages resulted in rapid host-range expansion of phage lysate. Lysis of non-permissive hosts PAK and PA14 was observed by round four. Clinical isolate strains CI00780, CI00795, and CI01788 were removed from the experiment after round three due to slow growth rates.

**Figure 3.**
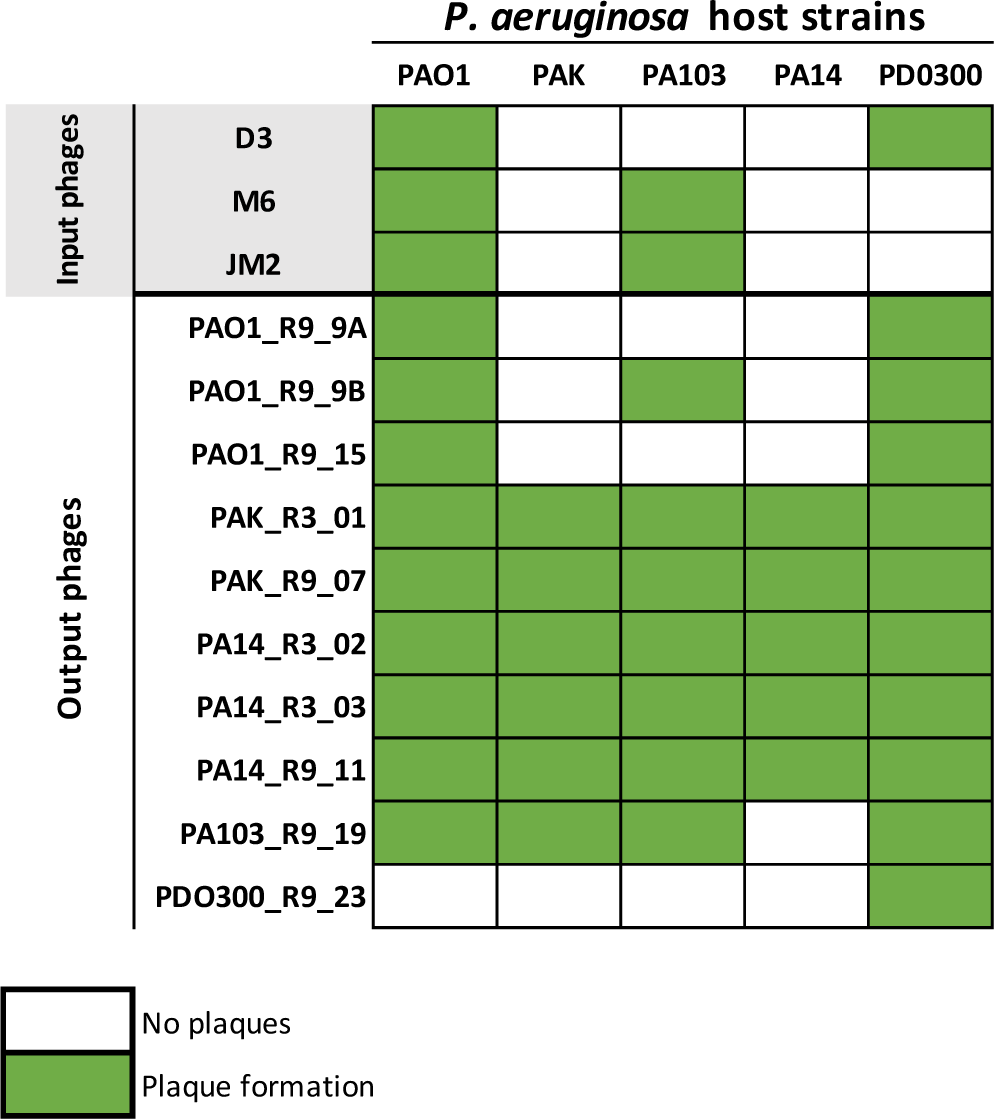
Host range of phages. Matrix representing host range of input phages and output phages isolated from Appelmans. Output phage naming scheme designates isolation host, isolation round, and phage number, respectively.

PA14_R3_03, PA14_R9_11) produced contigs between 37kbp and 39kbp, none of which were derived from input phages. Thus, we hypothesized that these output phage isolates may be derived from prophages originating from one of the bacterial hosts used in Appelmans.

### Intraspecies antagonism assays revealed presence of prophage in two host strains

To test the hypothesis that output phages may be derived from prophage, we conducted intraspecies antagonism assays between hosts used in Appelmans. All eight *P. aeruginosa* strains originally used in the experiment were cultured and subjected to induction by ultra-violet light and mitomycin C. Interspecies antagonism was measured by removing cells from these cultures and spotting the supernatants on lawns of all eight P. aeruginosa strains. Plaque formation was observed from the supernatant of strains PA103 and clinical isolate CI00795 (**Figure 4**). We also observed cleared zones of inhibition from six out of eight strains that is characteristic of tailocin activity (aka pyocins in *Pseudomonas*)(22, 23). Exposure of cultures to inducing agents did not produce higher levels of plaque formation or zones of inhibition compared to unexposed cultures, suggesting that these prophages are spontaneously produced from the hosts at low levels. Supernatant from PA103 could only form plaques on strain PAO1, and plaque formation was eradicated when the sample was exposed to chloroform. The prophage from PA103 may also be induced by exposure to heat, as lawns made from this host were riddled with plaques.

**Figure 4.**
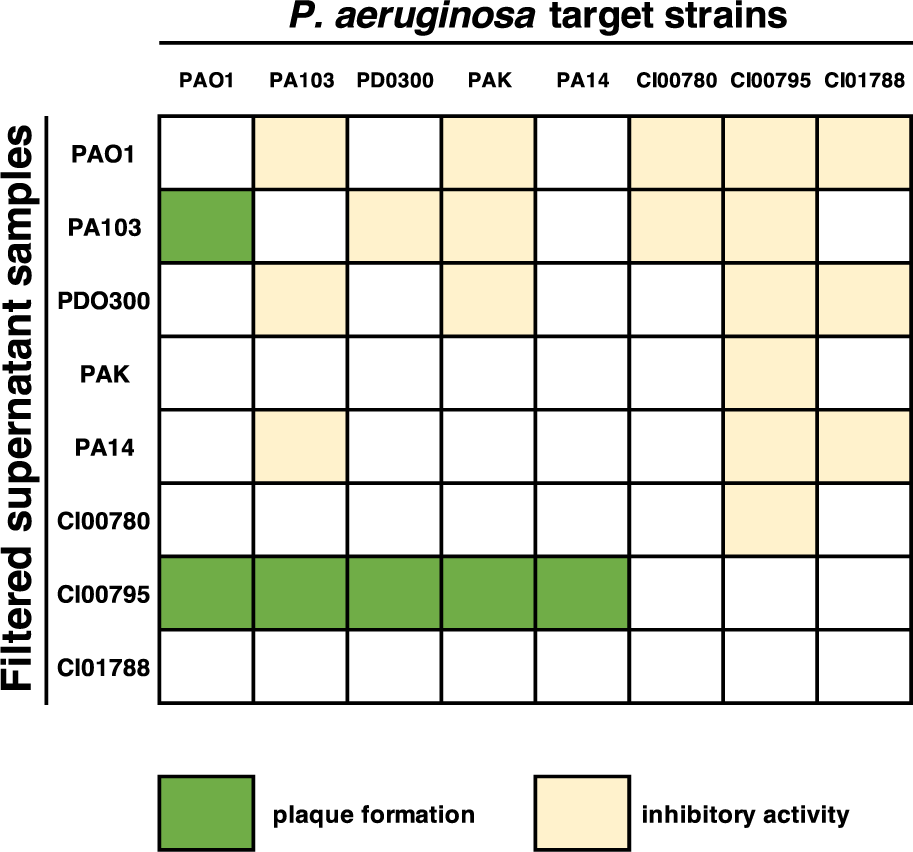
Intraspecies antagonism assay showed presence of active prophage in strains PA103 and CI00795. Inhibitory activity was observed from six out of eight hosts.

**Figure 4.**
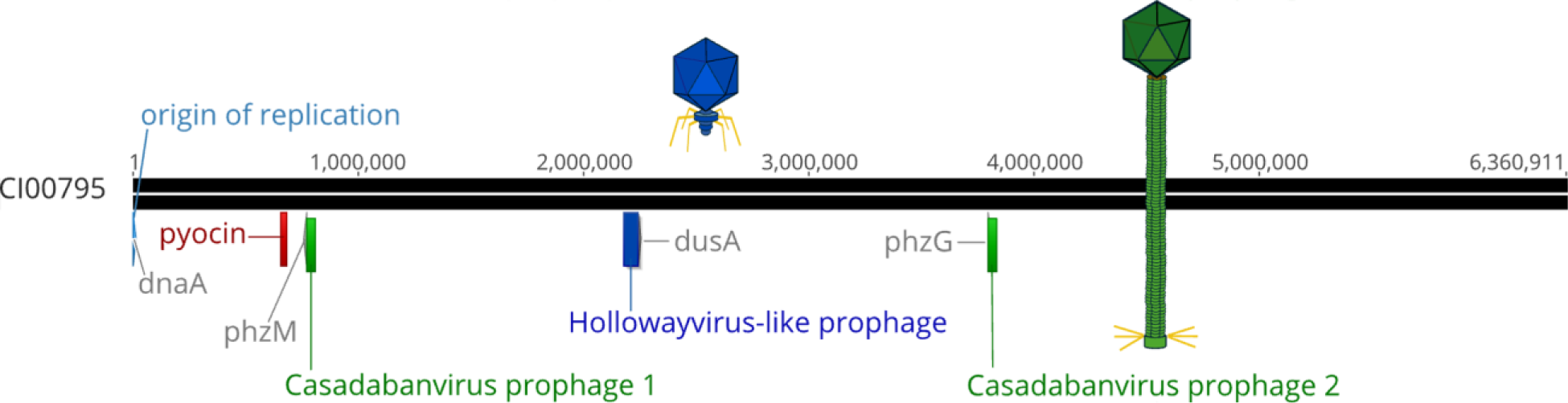
Genome map of *Pseudomonas aeruginosa* strain CI00795. Genome is oriented to beging with the origin of replication. The *Casadabanvirus* prophage is integrated at two locations: once near pyocyanin biosynthetic protein (phzM) and again near a phenazine biosynthesis FMN dependent-oxidase (phzG). The *Hollowayvirus* prophage is integrated near tRNA dihydrouridine synthase (dusA). A pyocin gene cluster is also present, located between anthranilate synthase component I (trpE) and anthranilate synthase component II (trpG).

Supernatant from strain CI00795 was able to form plaques on all five laboratory strains, including the two non-permissive hosts PAK and PA14, and was not susceptible to chloroform. Considering the narrow host range of the PA103 prophage, and that chloroform was regularly used to harvest and isolate phages throughout the Appelmans experiment, we determined it unlikely that the PA103 prophage contributed to host-range expansion.

Thus, we selected clinical isolate strain CI00795 for further analysis and sequencing.

### Five output phages are derived from a Casadabanvirus-like prophage in strain CI00795

To determine the origin of the five phage isolates with expanded host range, whole genome sequencing and hybrid assembly of the clinical host strain CI00795 genome was performed. Hybrid assembly using both short and long read data resulted in a single, circular contig of 6,360,911 bp (NCBI accession: CP158022, coverage: 234.7x). Genomic analysis of the CI00795 genome revealed that this host carries two prophages, one that is approximately 70kbp, and a second that is 37kbp (**Figure 4**). The 70kbp prophage is a *Hollowayvirus*-like prophage and is integrated between genes annotated as an exo-alpha-sialidase and a *tRNA dihydrouridine synthase* (dusA) (nucleotide position 2,176,677 -> 2,240,360). The 37kbp prophage is represented twice in the host genome, integrated in two locations: once between the genes phzM and hxlR (nucleotide positions 771,168 -> 808,371, henceforth referred to as “phi_1”), and again between phzG and a ORF annotated as a DNA excision repair complex subunit (nucleotide position 3,795,192 -> 3,832,395, henceforth referred to as “phi_2”). Sequence alignments confirmed that the five output phage samples were derived from the 37kbp prophage.

Further genomic analysis of the CI00795 prophages phi_1 and phi_2 indicated that this phage belongs to the genus *Casadabanvirus* (**Figure 5**). Phages in this genus, such as JDB24, have linear dsDNA genomes and are reported to be temperate, transposable, transducing, phages that, upon lysogenizing a host, can enhance the plaque formation of other infecting phages that are sensitive to CRISPR(18). They also integrate into multiple locations by transposition and can package up to 2kbp of heterogenous host DNA on the ends of their genome(24). Extraction and comparative genomics of phi_1 and phi_2 from the genome of CI00795 revealed that they are approximately 37,204 kbp and vary by 6 nucleotides (**Figure 6, Supplemental Table 1**). Variant analysis of the five output phages derived from the *Casadabanvirus* prophage revealed that four of the samples shared 100% identity with each other (excluding heterogenous host DNA regions) (**Supplemental Figure 3**), with the fifth sample sharing 99.99% identity. Only two unique mutations were identified in the five output phages that were not present in the two integrated prophages phi_1 and phi_2. All five output phages housed an intergenic insertion of “G” in a 5-nucleotide homopolymeric tract upstream of the tape measure protein. The fifth sample, PA14_R9_11, housed a nonsynonymous mutation in a putative tail fiber or baseplate protein.

**Figure 5.**
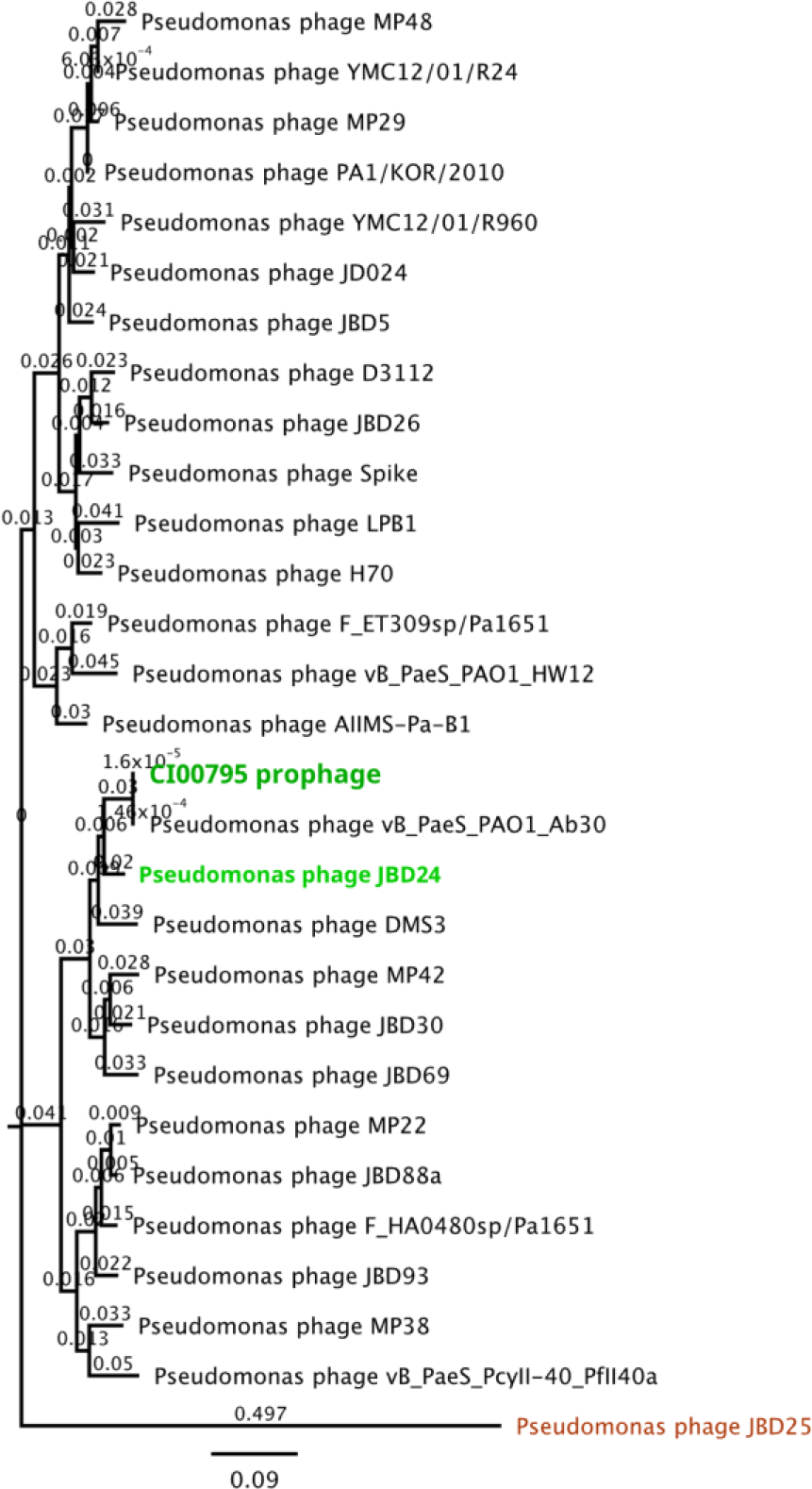
Phylogenetic analysis of the CI00795 37kbp prophage within the *Casadabanvirus* genus. Rooted neighbor-joining tree based on whole genome nucleotide multiple sequence alignment.; outgroup: JBD25. Branches represent substitutions per site.

### Host-range expansion of the Casadabanvirus prophage did not occur

Having established the origin of the five output phage samples, we sought to compare their host range with the initial host range of the *Casadabanvirus* prophage. To do this, we used PCR to distinguish the host range of the *Hollowayvirus* and *Casadabanvirus* prophages present in strain CI00795. We designed primers specific to the large terminase subunit of each prophage. We then spotted lysate from CI00795 onto lawns of the strains used in Appelmans and noted plaque formation. Plaques were harvested from each strain and used as the template DNA for PCR. This approach allowed us to confirm that the *Casadabanvirus* prophage can form plaques on host strains PAO1, PA103, PA14, PAK, and PDO300 (**Supplemental Table 2**). We also established that the *Hollowayvirus* prophage can form plaques on PAK and PDO300, but no positive results were observed for plaques harvested from the other strains. These results indicate that although the *Casadabanvirus* prophage was subjected to directed evolution conditions, it did not alter host range of the prophage for the host strains tested, as both the wild type and evolved prophages could infect all five bacterial strains.

## DISCUSSION

> *“In theory, there is no difference between theory and practice, while in practice there is.”* - Benjamin Brewster(25)

The directed evolution of phages involves methods aimed at promoting genetic and phenotypic variation to procure phages with desired characteristics. In theory, input phages propagated on input hosts will result in evolved input phages with the desired trait, and these dynamics can be modeled, and outcomes predicted(1). However, in practice, directed evolution studies can lead to variable and unexpected outcomes due to the complex dynamics of phage-host interactions and experimental conditions.

Prophages, dormant viral genomes housed within bacterial hosts, can influence these outcomes as an unexpected input and contribute to misinterpretation of results when not taken into consideration. This phenomenon poses challenges for some applications while offering solutions for others. For instance, in phage-host systems where lytic phages are scarce, the presence and induction of prophages can be viewed as a resource of potential therapeutic options. However, unwanted prophage content complicates the isolation of evolved phages and may impact regulatory approval of phages for therapeutic applications.

Here, we showed that induction of a prophage occurred during directed evolution in the *P. aeruginosa* phage-host system. The presence of this prophage was responsible for the observed host-range expansion of the pooled phage lysate. This prophage had a broader host range then the input phages, which allowed for it to persist throughout the experiment and become the most commonly isolated phage, even though the host it originated from was removed from the experiment after round three. Although the prophage was subjected to directed evolution conditions, it does not appear to have undergone host-range expansion. Although the induction and proliferation of prophage under directed evolution conditions was unexpected, it is not necessarily a surprising result given the nature and prevalence of prophages.

The number of prokaryotic genomes available has increased exponentially over the last two decades(26). This coupled with the ever-growing number of prophage prediction programs has allowed for the identification of prophage gene clusters in many bacterial species. Prophages are highly prevalent in human pathogens in particular with some strains harboring multiple prophages(7, 27–30). Furthermore, recent studies focused on the directed evolution of phages reported the induction of prophages. In the *Acinetobacter baumanii* phage-host system, prophages were observed to evolve via recombination which resulted in a prophage with expanded host range(17). In the *Klebsiella* phage-host system prophage induction was only observed when cultures were exposed to all three input phages, but not when cultures were exposed to individual phages(31). These findings, along with those of the current study, provide three examples of induction of prophage from pathogenic bacterial hosts during directed evolution.

Prophage induction and proliferation under directed evolution conditions is a fascinating, albeit mildly inconvenient, phenomenon; it presents an interesting problem for some, but a potential solution for others. For example, in some phage-host systems, strictly lytic phages are scarce, and only temperate phages are available for biocontrol or therapeutic applications(32, 33). If directed evolution methods can create conditions that allow for the induction and recombination of prophages, this may be a useful approach to procure evolved temperate phages with the desired host range. Although the use of temperate phages for phage therapy has been controversial, this approach is no longer out of reach in light of advancements in genetic engineering(34–36). Directed evolution methods, such as Appelmans, may present an opportunity to further study the evolution, induction pathways, and phage-host dynamics of prophages and temperate phages that could expand the pool of phages available for therapeutic applications.

In the current work, the prophage induced from a clinical isolate strain during Appelmans would be considered an interesting problem, as *Casadabanvirus* phages are known transposable, transducing, lysogenic phages that would not be suitable for phage therapy applications. Considering the prevalence of prophages in clinical isolates, and that directed evolution is commonly applied to target strains of clinical significance, the issue of prophage content is one that warrants greater attention. This raises the question: how does one solve the problem of unwanted prophage content? It is clear that sequencing input phages and evolved output phages is the most reliable method of ensuring that evolved phage isolates are derived from input phages, rather than being derived from prophages. Otherwise, the genetic origin of “evolved” phages isolated from directed evolution will remain unknown(2). The sequencing of bacterial strains is also important, particularly for those strains used for phage propagation and phage isolation. Having complete genome sequences for host is extremely useful in understanding resident phage related elements that could influence Appelmans results, such as tailocins and prophages. However, sequencing all bacterial target strains is not practical or feasible for many due resource constraints or time.

Here, we were able to utilize intraspecies antagonism assays of bacterial hosts to screen for potential prophage and tailocin content. This is a useful approach to establish a baseline of cross-strain inhibitory activity, including host range of resident prophage.

Conducting this assay prior to implementing Appelmans may also be helpful in selecting a suitable panel of propagation and target host strains. This data is also useful for interpretation of phage host-range testing of putatively evolved phages. Another approach could be to run a phage-free control of Appelmans to see if lysis occurs due to induction of tailocins or resident prophage. Here, we were also able to use PCR to efficiently screen plaques to identify and determine the host range of resident prophages. Alternatively, primers could be designed that are specific to input phages, and PCR used to quickly correlate observed plaque formation of pooled phage lysate with phage of origin, to avoid pursuing unwanted prophage derived isolates and preserve sequencing resources.

In conclusion, the presence and proliferation of prophages during directed evolution experiments presents both obstacles and opportunities in the quest to procure phages with desired characteristics for therapeutic applications. Such studies deepen our understanding of fundamental phage-host interactions. Understanding and managing the presence and influence of prophages in directed evolution studies is important for experimental design, allocation of resources, proper interpretation of results, and increasing the likelihood of favorable outcomes. Strategies such as sequencing input and output phages, sequencing hosts, performing intraspecies antagonism assays, and utilizing PCR for phage identification can aid in addressing these challenges.

## METHODS

### Bacterial and bacteriophage strains

All bacterial and bacteriophage strains are listed in **Supplemental Table 3**. All bacteria and phage were maintained at −80°C in 750 uL of LB media with 30% (wt/vol) glycerol. Bacterial strains were prepared for use by streaking for isolated colonies on 1.5% (wt/vol) LB agar plates and incubating at 37°C. Single colonies were used to prepare overnight cultures by inoculating 3mL of LB media and incubating shaking at 37°C. All input phages were propagated on host strain PAO1 Seattle. Host range of input phages was determined by spot plating 10uL of serially diluted phage stock onto lawns of bacterial hosts prepared on 3mL LB top agar (0.7% wt/vol) followed by incubation of plates at 37°C overnight and observation of plaque formation.

### Appelmans method

The Appelmans protocol is a method for passaging a phage cocktail on multiple hosts simultaneously typically in a 96-well plate(2). The experiment was set up as previously described by Burrowes et al.(2). Briefly, bacterial hosts were arranged in a 96-well plate by row, and evenly distributed across columns by adding 1ul of culture to 100uL of double strength LB media (**Figure 1**). The input phage cocktail contained a 1:1:1 ratio of phages D3, M6, and JM2, with a total input titer of 1×10^7^ pfu/mL. Ten-fold serial dilutions of the phage cocktail were made in LB media, and 100uL of each dilution was added to wells across columns, for a final volume of ∼200uL. The plate was then incubated overnight at 37⁰C and 200rpm. After overnight incubation, lysis of wells was observed and documented. All lysed wells and the first un-lysed or turbid well were collected in a 5mL falcon tube and chloroformed with 1:10 (v:v) ChCl3. The pooled phage lysate was centrifuged for 10 minutes at 10,000xg, and the supernatant was transferred and used as the input phage cocktail for the following round, for a total of nine rounds.

### Isolation of phages from Appelmans

Pooled phage lysate from Appelmans was serially diluted and plated on lawns of *P. aeruginosa* after round three and round nine. Resulting plaques from Appelmans protocol were double plaque purified by plating dilutions of the cocktail on each host individually using the top agar overlay method (overnight host culture and phage lysate dilution suspended in 3ml of LB top agar (0.7% w/v)). A plug of agar in the center of an individual plaque was taken and resuspended in a microcentrifuge tube containing 750ul LB media and 50ul ChCl3. This tube was then vortexed and spun down at 10,000xg for 10 minutes before the supernatant was collected. This process was repeated with the single isolated plaque supernatant to form a double isolated plaque supernatant. This 2x isolated plaque in LB media was then used as the working stock for subsequent host-range assays.

### Host-Range analysis of phages isolated from Appelmans

Host range of output phages was determined by observation of plaque formation on indicator lawns of bacteria; plaque formation indicates productive phage infection. To visualize plaques, 100ul of bacterial host culture was plated using the LB top agar overlay method to create a bacterial lawn. The top agar was allowed to set for 5-10 minutes before 2ul of phage dilutions were spotted across the lawn. Using a flame sterilized tungsten tool, each spot was streaked horizontally across the plate, flaming the tool between each use. After streaking, plates were incubated for 4-5 hours at 37⁰C and then left at room temperature for ∼8 hours before analysis. Streaks that resulted in individual plaque formation were considered positive.

### Bacterial and Phage genomic DNA extraction and sequencing

Bacterial gDNA for strain CI00795 was extracted using QIAgen DNeasy UltraClean Microbial kit as per manufacturer instructions. Phage stocks were amplified to a titer of ∼1×10^10^ PFU/mL by plating 1×10^5^ PFU/mL with 100ul of PAO1 Seattle overnight culture by LB top agar overlay. After overnight incubation at 37⁰C, the top layer of the cleared plate was collected in a 5mL microcentrifuge tube. After addition of 1:10 (v:v) ChCl3, 1.5mL of LB media, the tube was spun down at 10,000xg for 10 minutes and 1mL of the supernatant was transferred to a separate 5mL microcentrifuge tube. Phage gDNA was extracted using the Norgen Phage DNA Extraction Kit (including the optional protease and second elution steps). Bacterial and phage gDNA stocks were stored at −20⁰C, then submitted for sequencing with SeqCenter (Pittsburg, PA). At least 2 million 150 bp paired-end reads were obtained for every isolate. Strain CI00795 was also submitted to Plasmidsaurus for bacterial gDNA extraction and sequencing using Oxford Nanopore Technology.

### Assembly, genomic and variant analysis of bacterial hosts and phage isolates

Illumina reads were trimmed using fastp (version 0.22.0)(37). Fastqc was used to check for read quality of Illumina reads (version v0.11.3)(38). Reads were assembled using Unicycler(39) (version v0.4.3); a hybrid assembly was constructed for bacterial isolate CI00795 using the trimmed Illumina dataset and the long read dataset from Plasmidsaurus. The bacterial genome was annotated using Bakta(40). Phage genomes were annotated using Rast(41). Assembly statistics were generated using minimap2(42) and samtools(43). Variant analysis was conducted using breseq (44) and pairwise and multiple sequence alignments. Assemblies were visualized, and pairwise and multiple sequence alignments were conducted using Geneious Prime (version 11.0.15+10)(45).

### Intraspecies antagonism assays

In order to establish a baseline of inhibitory activity between *Pseudomonas* strains used in Appelmans and screen for resident prophage, instraspecies antagonism assays were conducted. Each bacterial host used in Appelmans was cross tested for intraspecies inhibitory activity (PAO1, PA103 PA14, PAK, PD0300, CI00780, CI00795, and CI01788). Bacterial cultures were grown and exposed to ultra-violet light, or mitomycin C (final concentration 1.0ug/mL) and incubated overnight; an uninduced culture was included as a control. The cultures were then centrifuged at 5,000 rpm for 15 minutes to pellet cellular debris, and supernatants were filter sterilized (0.2um). Filter sterilized samples were then ten-fold serially diluted and spotted at a volume of 10uL onto lawns of all bacterial hosts included in Appelmans. Plates were then incubated at 37⁰C overnight and observed for inhibitory activity and plaque formation.

### CI00795 prophage host-range determination by PCR

In order to determine if observed plaque formation on the five laboratory strains was due to the *Casdabanvirus* prophage or the *Hollowayvirus* prophage of strain CI00795, PCR primers were designed to target the large terminase subunit of each prophage. The forward and reverse primer sequences used to target the *Casdabanvirus* prophage in this study were 5’ CTAGCGTTGGTTAGAAGCCA 3’, and 5’ CGCCAGTGTCAAAAGAATCG 3’, respectively. The forward and reverse primer sequences for the *Hollowayvirus* prophage were 5’ CGAGTGACCACCTTCGTC 3’ and 5’ CTCACAGTCGCCAGTCAG 3’, respectively (primers were ordered from Invitrogen). To determine the host range of each prophage, 100uL of supernatant from strain CI00795 was added to 3mL of molten LB top agar, along with 100uL of overnight culture for each of the five *P. aeruginosa* laboratory hosts. Plaques were harvested from each bacterial lawn by picking a plaque using a 1000uL pipette tip and resuspending the plaque in 50uL of water. Samples were vortexed and allowed to sit at room temperature for 15 minutes. Samples were then placed in a heat block at 98°C for 5 minutes (46), then vortexed, and centrifuged at 5000xg for 5 minutes. Samples were then used as DNA template for PCR reactions using standard *Taq* DNA polymerase (NEB).

## Data availability

Raw read sequencing data for clinical isolate CI00795, input phages D3, M6, and JM2, and ten output phages are available under NCBI BioProject accession number PRJNA1104273. The assemblies for CI00795 and input phages D3, M6, and JM2 are available under the GenBank accessions CP158022, PP944329, PP944330, PP944331, respectively.

## ACKNOWLEDGEMENTS

We are grateful to James Bull for his feedback and riveting discussion regarding this manuscript. We would also like to acknowledge Jack Milstein for his preliminary data and technical support.

Research reported in this publication was supported by the National Institute of General Medical Sciences of the National Institutes of Health under Award Numbers P20GM104420, 3P20GM104420-09S1, and GM076040. The content is solely the responsibility of the authors and does not necessarily represent the official views of the National Institutes of Health. JS was supported by the University of Idaho Office of Undergraduate Research; ES was supported in part by R01GM076040.

